# Orbitofrontal Cortex is necessary for the behavioural expression, but not learning, of Pavlovian conditioned inhibition

**DOI:** 10.1101/2022.09.29.510151

**Authors:** Marios C. Panayi, Simon Killcross

## Abstract

Orbitofrontal cortex (OFC) lesions cause deficits in flexible behavioural control, most notably response inhibition and has historically been linked to theories of response inhibition. This general inhibition hypothesis of OFC function has since been rejected by evidence that inhibitory behavioural control can be expressed following OFC damage, however the functional role of the OFC in the explicit learning of conditioned inhibition remains untested. Here we test whether muscimol disruption of OFC function during the learning stage of a Pavlovian conditioned inhibition procedure disrupted the learning of conditioned inhibitory associations. Muscimol abolished Inhibitory behavioural control during the learning phase, however learning about the conditioned inhibitor was intact when tested drug free in subsequent summation and retardation tests of conditioned inhibition. Muscimol also significantly impaired acquisition to control cues whose cue-outcome relationship did not change. In a second experiment, conditioned inhibition was found not to play a significant role in cue extinction (non-reinforcement), an effect that was disrupted by intra-OFC infusion of muscimol. These results confirm that the OFC is not functionally necessary for the learning of inhibitory associations but is critical to both the enhancement and suppression of responding when environmental contingencies change.

---

Environmental cues can reliably signal motivationally significant outcomes allowing them to be predicted and to inform appropriate behaviours. Neural activity in the orbitofrontal cortex (OFC) tracks the learning and behaviour involved in this process. OFC neurons increase firing in the presence of reward predictive cues (Schoenbaum, Roesch, Stalnaker, & Takahashi, 2009; van Wingerden, Vinck, Lankelma, & Pennartz, 2010), suggesting that the OFC may form part of the representation of expected outcomes (Lucantonio et al., 2015; Rudebeck & Murray, 2014; Schoenbaum & Esber, 2010; Walton, Chau, & Kennerley, 2015). The OFC has been hypothesised as a site that integrates sensory and motivational information to adaptively increase or inhibit behaviour based on the up-to-the-moment expected value of predicted rewards (Rudebeck & Murray, 2014; Walton et al., 2015).

Surprisingly, disruption of OFC function does not disturb initial learning about cues that predict outcomes (Gallagher, McMahan, & Schoenbaum, 1999; McDannald, Lucantonio, Burke, Niv, & Schoenbaum, 2011) which is thought to depend on prediction-error signals that represent the discrepancy between expected and actual outcomes (Rescorla & Wagner, 1972; Schultz, 1998). Instead, current hypotheses attribute OFC involvement to situations in which initial learning must be modified by changes in the value or likelihood of the reward. For example, in extinction procedures when an expected reward is no longer delivered the expected value of the cue should be updated to reflect this new state of affairs, a process that is predicted to involve OFC function (Panayi & Killcross, 2014; Wilson, Takahashi, Schoenbaum, & Niv, 2014). In line with this prediction, Panayi & Killcross (2014) found that selectively inactivating rodent OFC function during extinction results in abnormally persistent responding to a cue that no longer predicts reward over many days.

While impaired extinction following OFC inactivation (Panayi & Killcross, 2014) is consistent with the hypothesis that the OFC is required to update behaviour based on the current value of predicted rewards, there are two alternative explanations of these results. One possibility is that the OFC is involved in the formation of inhibitory associations between events and expected outcomes, and therefore the rats never learn to inhibit their established behaviour. Historically, inhibitory explanations of OFC function have been ruled out by evidence that subjects with OFC damage can learn to inhibit a response if it has not been learned already (Murray, O’Doherty, & Schoenbaum, 2007; Rudebeck, Saunders, Prescott, Chau, & Murray, 2013). However, suppression of behaviour can occur via a number of alternative mechanisms that do not involve inhibition per se, such as behavioural competition, attention, and habituation (Panayi & Killcross, 2014; Rudebeck et al., 2013). The objective of this work is to provide the first direct test of the role of the OFC in the acquisition of conditioned inhibitory associations.

Extinction learning has been argued to involve predominantly new context-dependent inhibitory learning rather than unlearning of the original association (Bouton, 1993; Delamater, 2004). Therefore, a second explanation of the role of the OFC in extinction learning is the formation of new state information to support this new inhibitory learning (Wilson et al., 2014). In the absence of this new state representation Wilson et al (2014) predict that the rate of extinction will be retarded since the original association will undergo unlearning rather than the formation of new context specific inhibitory learning. Importantly, the OFC is proposed to only be involved in the representation of task states when the states are not explicitly signalled but instead require the use of memory to infer a new “latent” state.

A Pavlovian conditioned inhibitor can be established by learning that a cue that is normally reinforced (A+) is non-reinforced in the presence of a second cue (AX-) (Papini & Bitterman, 1993; Rescorla, 1969). This resembles an extinction procedure except that non-reinforcement is explicitly signalled by a second cue (X). We hypothesise that if the OFC is necessary for establishing or learning about latent task states (Wilson et al., 2014), OFC inactivation should not interfere with learning about an explicitly signalled conditioned inhibitor. However, if the OFC is simply involved in the acquisition of inhibitory associations, then OFC inactivation should disrupt the expression and subsequent learning of conditioned inhibition.

## Materials and Methods

### Animals

Fifty-six adult male Long-Evans rats (302-406 g prior to surgery; Monash Animal Services, Gippsland, Victoria, Australia) were used (experiment 1, N = 32; experiment 2, N = 24). Rats were housed four per cage in ventilated Plexiglass cages in a temperature regulated (22 ± 1°C) and light regulated (12h light/dark cycle, lights on at 7:00 AM) colony room. At least one week prior to behavioural testing, feeding was restricted to ensure that weight was approximately 95% of ad libitum feeding weight, and never dropped below 85%. All animal research was carried out in accordance with the National Institute of Health Guide for the Care and Use of Laboratories Animals (NIH publications No. 80-23, revised 1996) and approved by the University of New South Wales Animal Care and Ethics Committee.

### Apparatus

Behavioural testing was conducted in eight identical operant chambers (30.5 × 32.5 × 29.5 cm; Med Associates) individually housed within ventilated sound attenuating cabinets. Each chamber was fitted with a 3-W house light that was centrally located at the top of the left-hand wall. Food pellets (45 mg dustless precision grain-based pellets; Bio-serv, Frenchtown, NJ, USA) could be delivered into a recessed magazine, centrally located at the bottom of the right-hand wall. The top of the magazine contained a white LED light that could serve as a visual stimulus. Access to the magazine was measured by infrared detectors at the mouth of the recess. Two retractable levers were located on either side of the magazine on the right-hand wall. A speaker located to the right of the house light could provide auditory stimuli to the chamber. In addition, a 5-Hz train of clicks produced by a heavy-duty relay placed outside the chamber at the back-right corner of the cabinet was used as an auditory stimulus. The chambers were wiped down with ethanol (80% v/v) between each session. A computer equipped with Med-PC software (Med Associates Inc., St. Albans, VT, USA) was used to control the experimental procedures and record data.

Locomotor activity was assessed in a set of 4 rat open field arenas (Med Associates Inc., St. Albans, VT, USA) individually housed in light and sound attenuating cabinets. A 3-W light attached on the top left corner of the sound attenuating cabinet provided general illumination in the chamber and was always on. A 28 V DC fan on the right-hand wall of the sound attenuating cabinet was also left on throughout testing to mask outside noise. The floor of the open field arena was smooth plastic and the four walls were clear Perspex with a clear Perspex roof containing ventilation holes. The internal dimensions of the chamber were 43.2 × 43.2 × 30.5 cm (length x width x height). Two opposing walls contained an array of 16 evenly spaced infrared detectors set 3 cm above the floor to detect animal locomotor activity. A second pair of infrared beam arrays was set 14 cm above the floor on the remaining walls to detect rearing behaviours. Infrared beam breaks were recorded using a computer equipped with Activity Monitor software (Med Associates Inc., St. Albans, VT, USA) which provided a measure of average distance travelled based on beam break information.

### Surgery

Surgery was performed prior to training in experiment 1 and after initial training in experiment 2. Bilateral guide cannulae were surgically implanted targeting the lateral OFC. Rats were anesthetized with isoflurane, their heads shaved, and placed in a stereotaxic frame (World Precision Instruments, Inc., Sarasota, FL, USA). The scalp was incised, and the skull exposed and adjusted to flat skull position. Two small holes were drilled for the cannulae using a high-speed drill, and four holes were hand drilled on different bone plates to hold fixing screws. Bilateral stainless steel guide cannulae (26 gauge, length 5mm below pedestal; Plastics One, Roanoke, VA, USA) were lowered into the lateral OFC (AP: +3.5 mm; ML: ±2.2 mm; D-V: -4.0 mm from bregma). Cannulae were held in place by dental cement and anchored to the skull with 4 fixing screws. Removable dummy cannulae were inserted into the guide cannulae to prevent them from blocking. After one week of postoperative recovery, rats were returned to food restriction for 2 days prior to further testing.

### Drugs and infusions

The GABA_A_ agonist muscimol (Sigma-Aldrich, Switzerland) was dissolved in 0.9% (w/v) non-pyrogenic saline to obtain a final concentration of 0.5 *μ*g/0.5 *μ*l. Non-pyrogenic saline 0.9% (w/v) was used as the saline control. During infusions, muscimol or saline was infused bilaterally into the lateral OFC by inserting a 33-gauge internal cannula into the guide cannula which extended 1 mm ventral to the guide tip. The internal cannula was connected to a 25 *μ*l glass syringe (Hamilton Company, Reno, NV, USA) attached to a microinfusion pump (World Precision Instruments, Inc., Sarasota, FL, USA). A total volume of 0.5 *μ*l was delivered to each side at a rate of 0.25 *μ*l/min. The internal cannula remained in place for an additional 1 min after the infusion and then removed. During the infusion procedure animals could move freely in a bucket to minimize stress. Dummy cannulae were removed prior to, and replaced immediately after, infusions. For the two training sessions prior to infusions, all animals received dummy infusions which were identical to the infusion procedure, but no liquids were infused. These dummy infusions were performed to familiarize the rats with the microinfusion procedure and thereby minimize stress. Similar handling without cannula insertion occurred prior to all behavioural testing sessions.

### Histology

Following completion of behavioural testing, rats received a lethal dose of sodium pentobarbital. Brains were extracted, frozen, and sectioned coronally at 40 *μ*m through the lateral OFC with a cryostat. Every third section was collected on a slide and stained with cresyl violet. The location of cannula tips was determined under a microscope by a trained observer, unaware of subject’s group designations, using the boundaries defined by the atlas of George Paxinos and Watson (1998). Rats with bilateral cannulae placements outside the lateral or dorsolateral OFC were excluded from statistical analyses.

### Magazine training

Prior to each experiment, all rats were familiarised with retrieving rewards from the magazine in a session of magazine training that lasted approximately 32 mins. Rewards, consisting of two pellets delivered 0.25s apart, were delivered randomly throughout the session every 120s until 32 pellets were delivered. The house light was kept illuminated throughout the session.

### General session parameters

All sessions consisted of a number of trials in which 10s auditory and/or visual cues (conditioned stimuli; CS) were presented. Visual cues designated as X and Y were flashing panel lights (0.1 s illuminated, 0.1s off) or extinguishing the house light (identity counter balanced). Visual cue Z was always a flashing magazine light (0.1 s illuminated, 0.1s off) for all animals. Auditory cues A and B were a 5 Hz train of clicks or a 78 dB white noise (identity counter balanced). Auditory cue C was always an 84 dB, 2.6 kHz tone. On rewarded trials (denoted by the symbol ‘+’) a single reward pellet was delivered upon CS termination. On non-reinforced trials (denoted by the symbol ‘-’), no reward was delivered. The variable inter-trial-interval was 90s (± 45s). Unless stated, only a single training session occurred per day and cue order was randomised. All animals were handled in the infusion bucket for 5 minutes prior to each session and handled similarly regardless of whether drug infusions were administered. This was done to equate handling cues and stress on all training days.

### General experimental design

Experiments 1 and 2 were designed to establish cue X as a conditioned inhibitor (Figure 1B and 1D). This was achieved by first training cue A as a valid predictor of reward (A+, Stage 1) before training the compound AX as a valid predictor of the absence of reward (AX-, Stage 2). Experiment 1 was designed such that A+ and AX- were trained as a discrimination within the same sessions (Figure 1B), a feature negative discrimination procedure that is commonly used to generate robust conditioned inhibition to cue X (Papini & Bitterman, 1993). Experiment 2 was designed such that AX- was presented in separate sessions only after A+ was well trained instead of within the same session (Figure 1D). This design has been used to probe the formation of conditioned inhibition in extinction procedures (Rescorla, 1979), and provided a test of whether the extinction parameters employed by (Panayi & Killcross, 2014) promoted the formation of conditioned inhibition.

**Figure 1.**
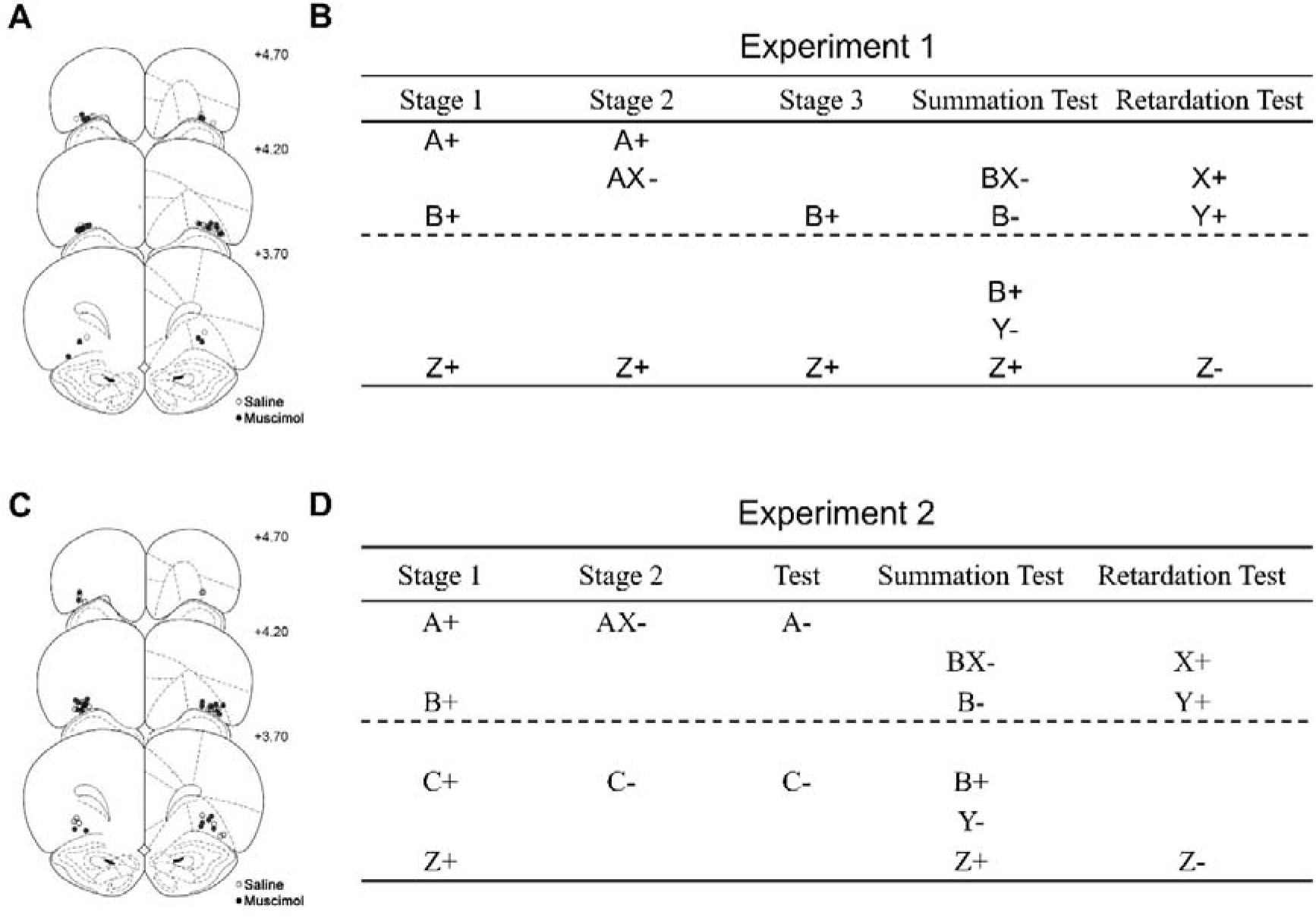
Experiment 1 **(A)** Schematic representation of cannulae tip placements in the OFC. Coronal sections are identified in mm relative to bregma (Paxinos and Watson, 1997). **(B)** The design of experiment 1 intended to establish cue X was a conditioned inhibitor. This is achieved by training cue A as a reliable predictor of reward unless it is simultaneously presented in compound with cue X. The main design is depicted above the dashed line, whereas additional control cues are depicted below the dashed line. A and B are auditory cues (white noise and click), C was always a tone, X and Y are visual cues (house light off and panel lights on), Z was always a flashing magazine light and the symbols “+” and “-” denote reward and non-reward respectively. Infusion of saline or muscimol occurred during stage 2. Experiment 2 **(C)** Schematic representation of cannulae tip placements in the OFC. **(D)** The design of experiment 2 intended to establish cue X as a conditioned inhibitor. This is achieved by training cue A as a reliable predictor of reward unless it is simultaneously presented in compound with cue X. The key aspects of the procedure are highlighted above the dashed line and additional control cues are presented below the dashed line. A and B are auditory cues (white noise and click), X and Y are visual cues (house light off and panel lights on), Z was always a flashing magazine light and the symbols “+” and “-” denote reward and non-reward respectively. Infusion of saline or muscimol occurred during stage 2.

Experiments 1 and 2 employed both a summation and a retardation test to assess whether cue X had become a conditioned inhibitor (Rescorla, 1969). Briefly, a cue can be considered to be a conditioned inhibitor if it is able to transfer its inhibitory properties to other predictive cues (summation test), and it should be harder to transform an inhibitory cue into an excitatory cue relative to a neutral cue (retardation test). In summation tests, cue X is presented in compound with another excitatory cue, BX, and if responding is inhibited/lower to this compound than to B alone, then cue X is considered to be a conditioned inhibitor. However, it is possible that lower responding to the compound BX in a summation test is caused by generalisation decrement i.e. a reduction in responding due to generalisation between compound AX and BX, or due to external inhibition caused by the novel pairing of cues B and X. Alternatively, in a retardation test, if X has accrued inhibitory properties, the rate of acquisition to cue X will be lower than acquisition to a novel. Unlike summation tests, there is no issue of generalization decrement in a retardation test, however slower acquisition to cue X may be caused by reductions in attention to cue X as it is presented repeatedly during AX-training (i.e. latent inhibition). A consistent attentional explanation of both the summation and retardation test is not possible. It has therefore been argued that to rule out alternative explanations, both summation and retardation tests are needed to establish that cue X has acquired conditioned inhibitory properties (Papini & Bitterman, 1993).

#### Experiment 1 acquisition days 1-4

During each acquisition session a total of 36 trials were presented consisting of 12 A+, 12 B+ and 12 Z+ trials. Each session lasted 60 mins. On days 3 and 4, all animals received dummy infusions immediately prior to the session.

#### Experiment 1 feature negative training days 5-10

During each feature negative training session all animals received a total of 36 trials consisting of 10 A+, 20 AX- and 6 Z+ trials. The non-rewarded AX-trials consisted of the simultaneous presentation of the audio-visual cues A and X. The A+/AX-feature negative discrimination was used to establish cue X as a conditioned inhibitor. Prior to each feature negative training session all animals received an infusion of either muscimol or saline targeting the OFC.

#### Experiment 1 cue retraining days 11-12

During cue retraining sessions a total of 36 trials were presented, consisting of 18 B+ and 18 Z+ trials. This retraining was done to ensure that responding to cue B was high prior to the summation test and to assess any persistent effects of the infusion procedure.

#### Experiment 1 summation probe test day 13

The summation probe test consisted of 27 trials (45 mins session length) in the following order: first 3 Z+ and 3 B+ trials (order: Z+, B+, B+, Z+, B+, Z+). This rewarded start ensured high responding to the critical target cue B. Then 2 B- and 2 BX-trials were presented, followed by a Z+ trial. This cycle of 5 trials (B-/BX-/Z+) was repeated 2 more times. The B-/BX-cues were probe trials to test whether cue X had acquired inhibitory properties that transferred to cue B. Rewarded Z+ trials were interspersed throughout the session to maintain responding throughout the probe trials. Finally, all animals received 6 presentations of Y- at the end of the session. This pre-exposure to cue Y was done to minimise any external inhibition that may occur during the retardation test that followed.

#### Experiment 1 retardation test days 14-16

The retardation test sessions contained 36 trials consisting of 12 X+, 12 Y+ and 12 Z-trials (order randomized). This test shows whether the prior inhibitory training with cue X impairs subsequent excitatory acquisition relative to the novel cue Y. The non-rewarded cue Z was designed to prevent animals from responding non-discriminatively to all cues during this acquisition session.

#### Experiment 1 consumption test days 17-18

Following the retardation test, all animals were given a consumption test to assess whether muscimol infusions into the OFC impaired the motivation or timing of pellet consumption, which may explain reduced performance during the Stage 2 feature negative training following infusions. On day 17, all animals were given a dummy infusion immediately prior to entering the test chamber. Prior to the session, 40 pellets were placed in the magazine. All animals were given 30 minutes in the chamber. Magazine behaviour was recorded during this session for analysis, but there were no programmed events throughout the session. On day 18 all animals were infused with muscimol or saline before being entered for a second consumption test identical to that on day 17.

#### Experiment 2 acquisition days 1-9

During acquisition sessions there were a total of 36 trials, consisting of 9 A+, 9 B+, 9 C+ and 9 Z+ trials. Each session lasted 60 mins. Animals were entered for 2 sessions per day for stage 1 training for a total of 12 sessions across days 1-6. Animals were then returned to free feeding and surgery was performed. Immediately following post-operative recovery all animals were returned to food restriction 2 days prior to further acquisition. Post-operative acquisition on days 7-9 were identical to pre-surgical Stage 1 acquisition, except that only a single session was administered per day. On the final two days all animals received dummy infusions immediately prior each session.

#### Experiment 2 feature negative training days 10-13

During the feature negative training, each session consisted of 36 trials such that there were 18 AX- and 18 C-trials. Infusions of saline or Muscimol were administered immediately to separate groups (matched on performance to all cues) prior to each of these sessions.

#### Experiment 2 extinction test day 14

During the extinction test there were a total of 24 trials consisting of 12 A- and 12 C-trials.

#### Experiment 2 summation and retardation tests

The summation and retardation tests were identical to those described in Experiment 1.

#### Experiment 2 Locomotor activity

At the end of training, animals were locomotor screened following an additional drug infusion (either saline or Muscimol) to evaluate whether drug infusions affected general activity levels between groups.

##### Data analysis

CS responding was operationalized as the number of magazine entries during the 10s CS. PreCS responding was operationalized as the frequency of responding during the 10s immediately preceding the 10s CS and was used as a measure of baseline responding to the testing context. All data were analysed with mixed ANOVAs, and significant interactions of interest were followed up with ANOVAs on the relevant subset of data. A Tukey correction was used when relevant simple effects were performed. Following significant omnibus ANOVA tests, planned linear orthogonal trend contrasts and their interactions between groups were also analysed to assess differences in rates of acquisition over days or blocks of trials when relevant.

For clarity, cue Z was analysed separately as it was a control cue that was not counterbalanced (a flashing light in the magazine) and elicited a different pattern of responding to the other cues.

## Results

### Experiment 1: OFC inactivation disrupts the expression but not the acquisition of conditioned inhibition

#### Histology

Cannulae placements are depicted in Figure 1A (representative image shown in Supplemental Figure S1A). Two animals were excluded from further analysis due to misplaced cannulae. During training a further two animals assigned to the saline group were excluded and were not trained further as they failed to acquire magazine training after several days. Final numbers for infusion groups in Experiment 1 were saline (n = 13) and muscimol (n = 15).

#### Baseline responding

Rates of baseline responding during the 10s PreCS period did not significantly differ between groups during any of the testing phases, and justified the analysis of CS-PreCS difference scores as measures of discriminative responding to the cues in subsequent analyses (data not shown). Briefly, Group x Day mixed ANOVAs were run separately for each stage of testing. During stage 1 acquisition (main effect of Group F(1, 26) = 3.20, *p* = .09; Group x Day interaction F(3, 78) = 1.47, *p* = .23), stage 2 feature negative training, stage 3 cue re-training, summation and retardation tests all Group and Group x Day interactions failed to reach significance (all F < 2.01, *p* > .16).

#### Stage 1: Acquisition (Days 1-4)

Prior to infusions, all rats acquired conditioned responding to cues A+ and B+ (Figure 1A; main effect of Day *F*(3,78) = 27.49, *p* < .001, linear trend *t*(26) = 7.63, *p* < .001, all remaining effects and interactions *F* < 1.34, *p* > 26).

#### Stage 2: OFC inactivation abolishes selective inhibition of behaviour (days 5-10)

The saline group successfully acquired the feature negative A+/AX-discrimination (Figure 2B; significant Cue*Day interaction *F*(5,60) = 6.23, *p* < .001, and main effects of Cue *F*(1,12) = 5.57, *p* = .036, and Day *F*(5,60) = 7.50, *p* < .001) by increasing responding over days to the rewarded cue A+ (significant linear increase *t*(12) = 6.39, *p* < .001), and selectively inhibiting responding to the non-rewarded compound AX- (non-significant linear trend *t*(12) = 0.56, *p* = .588). In contrast, the muscimol group did not significantly discriminate between A+ and AX- (non-significant Cue*Day interaction *F*(5,70) = 1.06, *p* = .388, or main effect of Cue *F*(1,14) = 0.01, *p* = .920, but did show a signficant main effect of Day *F*(5,70) = 3.65, *p* = .005, and a linear increase over days *t*(14) = 2.75, *p* = .016). These group differences were supported by a significant Group*Cue*Day interaction (*F*(5,130) = 4.91, *p* < .001, as well as a significant Group*Day interaction *F*(5,130) = 2.94, *p* = .015, and main effect of Group *F*(1,26) = 8.69, *p* = .007). Therefore, the saline group showed behavioural evidence of selective inhibition during the A+/AX-feature negative discrimination which was abolished by intra-OFC infusions of muscimol. While this suggests that OFC function is necessary for selective inhibitory control of behaviour, it is unclear whether learning about the conditioned inhibitor X was also impaired.

**Figure 2.**
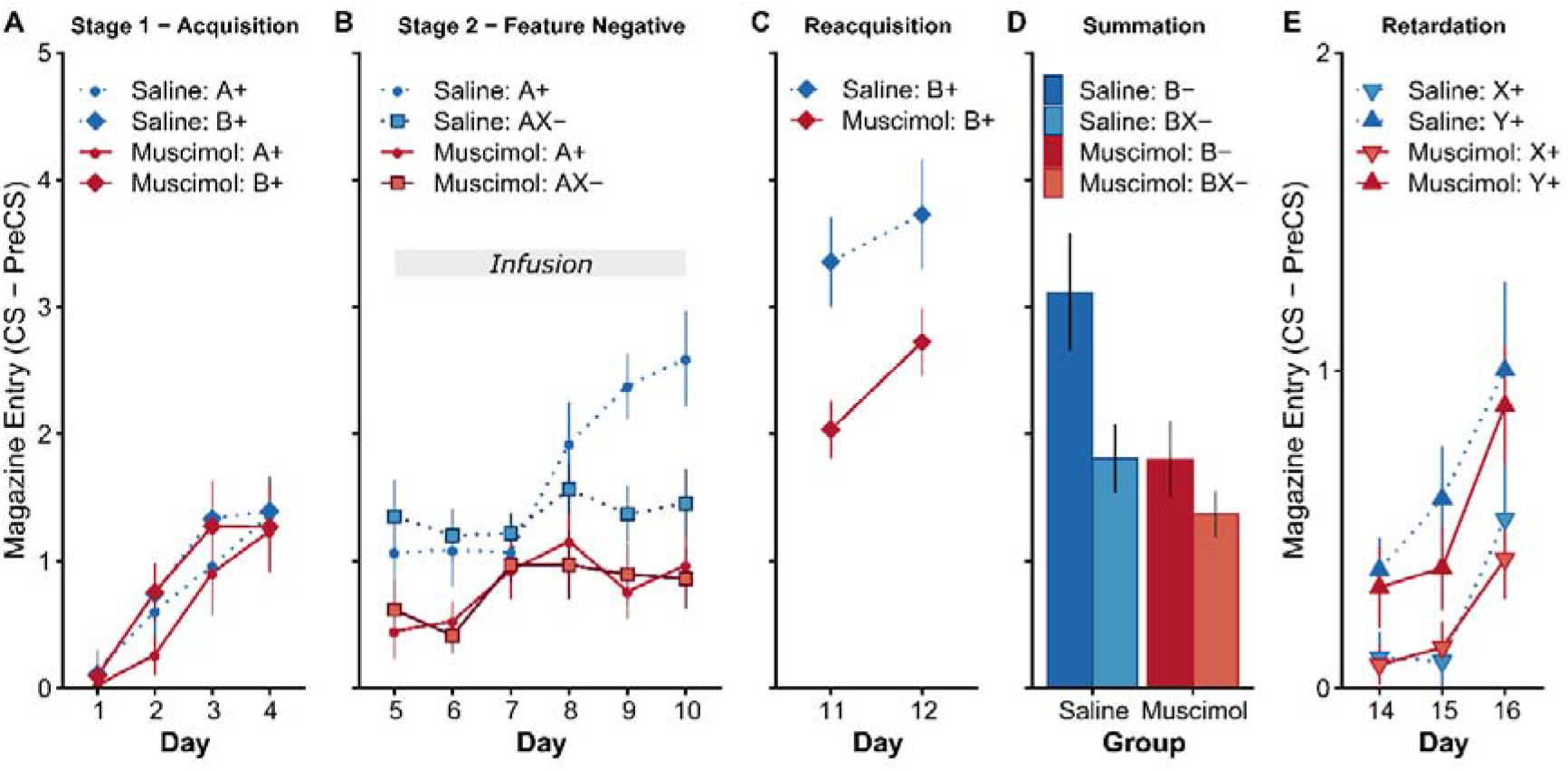
OFC is necessary for the expression but not the learning of conditioned inhibition. Rates of discriminative magazine responding in Experiment 1 presented as CS-PreCS difference scores in 10s. **(A)** Acquisition of responding to cues A+ and B+ in Stage 1. **(B)** Acquisition of the A+/AX-discrimination following saline or muscimol infusions. Following saline infusions, responding to A+ was greater than AX-whereas muscimol infusions abolished differences in responding to A+ and AX-. **(C)** Reacquisition to control cue B+ in stage 3 in the absence of infusions revealed significantly lower responding in the muscimol group. **(D)** A summation test revealed lower responding to BX- than B- in both the saline and muscimol groups (see Supplemental Figure S2 for first 2 trials of the summation test). **(E)** A retardation test revealed significantly lower responding to X+ than to the novel control cue Y+ in both the saline and muscimol groups. Error bars depict ±SEM.

#### Stage 3: Retraining (days 11-12)

In stage 3, the overall suppression of responding observed in stage 2 persisted temporarily during retraining to cue B drug-free (Figure 2C; significant main effect of Group *F*(1,26) = 7.95, *p* = .009, and Day *F*(1,26) = 7.81, *p* = .010, but no Group*Day interaction *F*(1,26) = 0.70, *p* = .411). This suggests that the effect of muscimol infusion in stage 2 temporarily, and non-selectively, lowered overall performance when trained drug free in stage 3. It is possible that this effect is due to the disruption of motivation for the reward or an overall suppression of motor function, however these possibilities were ruled out by the results of the consumption test administered at the end of testing (reported below).

#### Summation and retardation test: OFC inactivation during training does not prevent the acquisition of conditioned inhibition (days 13-16)

While OFC inactivation successfully abolished the expression of selective conditioned inhibition in the feature negative discrimination (stage 2, Figure 2C), it is not clear whether this indicates a failure of acquisition of conditioned inhibition or just impaired behavioural expression. To address this question directly, summation and retardation tests of conditioned inhibition (Papini & Bitterman, 1993; Rescorla, 1969) were administered drug-free to allow for any latent learning to be expressed.

The results of the summation test (Figure 2D) revealed that both groups respond less to the compound BX- than B- (significant main effect of Cue *F*(1,26) = 7.78, *p* = .010) which suggests that cue X successfully acquired inhibitory properties during the feature negative training (stage 2). The muscimol group had lower overall responding to all cues than the saline group (main effect of Group *F*(1,26) = 8.26, *p* = .008), but the magnitude of the summation effect was not significantly different between groups (Group*Cue interaction *F*(1,26) = 1.96, *p* = .173). While the plotted data might suggest that the summation effect was weaker in the muscimol group, closer inspection of the data revealed that this was due to the overall lower levels of responding in the muscimol group and more rapid extinction of behaviour during the non-reinforced trials see (Supplemental Figure S2 for data from the first block of non-reinforced trials during summation test). These findings suggest that intra-OFC infusions of muscimol did not disrupt the acquisition of conditioned inhibition to cue X as assessed by a summation test. However, a reduction in responding to the BX compound could also be explained by enhanced attention to cue X, generalisation decrement, or external inhibition.

To rule out these alternative explanations a retardation test was conducted in which the rate of acquisition to X+ was compared to the relatively novel cue Y+. If cue X has acquired inhibitory properties, then acquisition should be slower to X+ than Y+. Importantly, this result would be incompatible with an account of the summation test appealing to enhanced attention to X, which would predict an increase in the rate of learning. During the retardation test (Figure 2E; days 14-16) acquisition to target cue X+ was significantly slower than to control cue Y+ in both groups (significant main effect of Cue *F*(1,26) = 24.78, *p* < .001, and Day *F*(2,52) = 12.56, *p* < .001, all remaining *F* < 1.08, *p* > .348). Retarded acquisition to X+ relative to Y+ suggests a significant retardation effect of similar magnitude in both the saline and muscimol groups. Together, the results of the summation and retardation tests demonstrate that cue X has indeed acquired conditioned inhibition, even though OFC inactivation abolished discriminative performance during the feature negative training learning phase in stage 2.

#### Control cue Z: OFC inactivation disrupts Pavlovian acquisition

The adverse consequences of disrupting OFC function are usually only detected when task contingencies change (Rudebeck & Murray, 2014; Wilson et al., 2014), but rarely during the initial acquisition of a task when contingencies and response requirements remain stable (Walton, Behrens, Noonan, & Rushworth, 2011). Therefore, it is surprising that OFC inactivation during the feature negative discrimination also disrupted the acquisition of responding to control cue Z (Figure 3). Prior to infusions there were no differences in acquisition to Z+ (Figure 3A; main effect of Group *F*(3,78) = 0.17, *p* = .916, Group*Day interaction *F*(3,78) = 0.17, *p* = .916, main effect of Day *F*(3,78) = 2.58, *p* = .059; significant linear increase in responding over days *t*(26) = 2.25, *p* = .033). However, during stage 2, infusion of muscimol significantly impaired acquisition (Figure 3B; significant main effect of Group *F*(1,26) = 13.32, *p* = .001, Group*Day interaction *F*(3,78) = 0.17, *p* = .916, and main effect of Day *F*(3,78) = 2.58, *p* = .059). There was evidence that the muscimol group did acquire responding over days 5-10 (significant linear increase over days, muscimol *t*(26) = 3.55, *p* = .002, saline *t*(26) = 6.43, *p* < .001), however while both groups had similar levels of responding on day 5 (*t*(26) = −0.28, *p* = .779), responding was significantly lower in the muscimol group on days 6-10 (Day 6 *t*(26) = −3.00, *p* = .006, Day 7 *t*(26) = −3.15, *p* = .004, Day 8 *t*(26) = −2.29, *p* = .031, Day 9 *t*(26) = −3.32, *p* = .003, Day 10 *t*(26) = −2.67, *p* = .013). This suggests that the OFC plays a role in Pavlovian acquisition.

**Figure 3.**
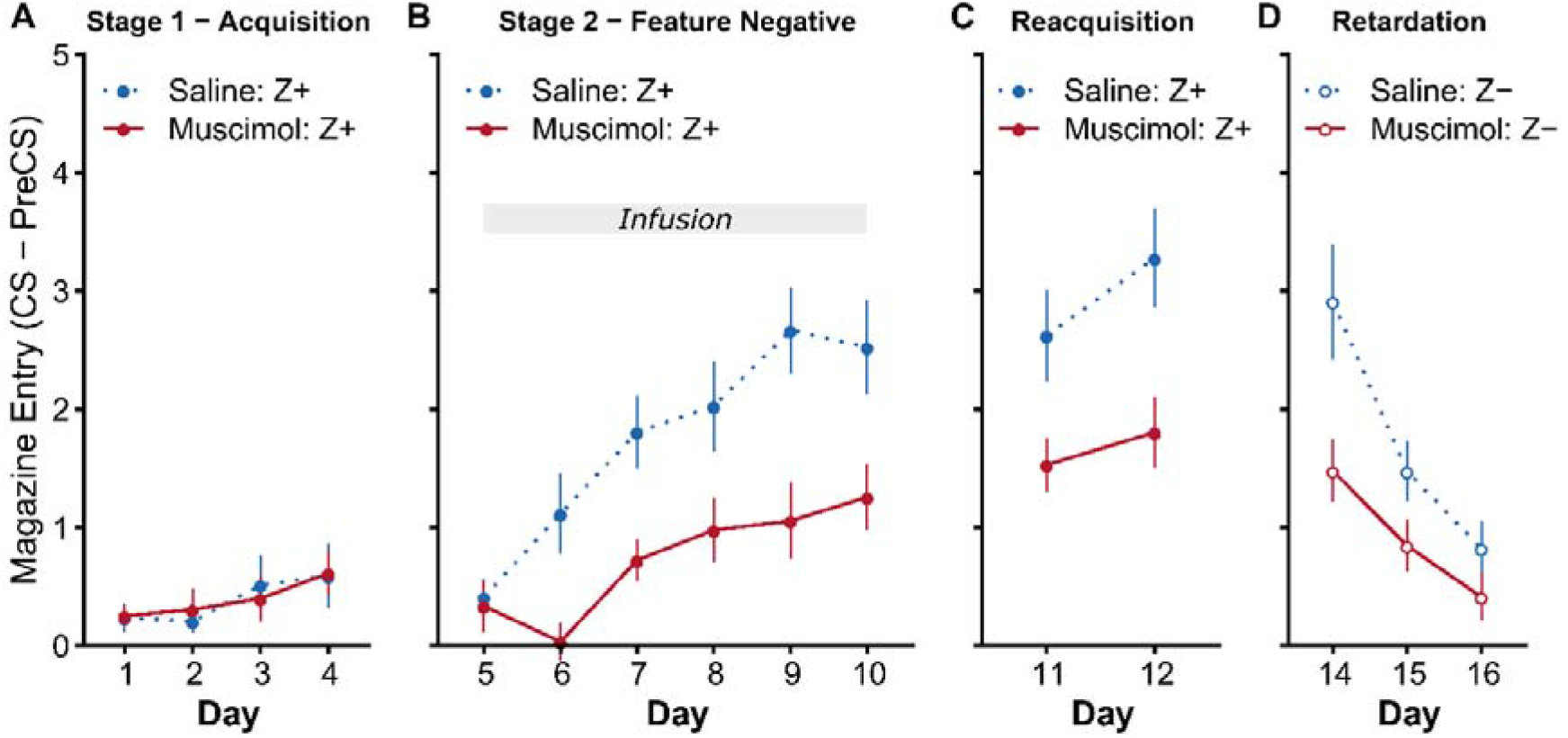
OFC inactivation disrupts Pavlovian acquisition. Responding to control cue Z in Experiment 1 in the test phases described in Figure 2. **(A)** There were no significant group differences in stage 1 acquisition prior to drug infusions. **(B)** The muscimol group responded significantly lower than the saline group in stage 2 and this difference persisted when tested drug free in **(C)** stage 3 and during **(D)** extinction of cue Z in the retardation test. Rates of discriminative magazine responding are presented as CS-PreCS difference scores in 10s. Error bars depict +SEM.

Suppressed responding to cue Z persisted in the muscimol group when trained drug-free in subsequent sessions (Figure 3C, days 11-12; significant main effect of Group *F*(1,26) = 7.85, *p* = .009, Day *F*(1,26) = 15.18, *p* = .001, but no Group*Day interaction *F*(1,26) = 2.54, *p* = .123). This difference was also evident when cue Z underwent extinction during the retardation test (Figure 3D, days 14-16; significant main effect of Group *F*(1,26) = 6.11, *p* = .020, Day *F*(2,52) = 32.34, *p* < .001, and Group*Day interaction *F*(2,52) = 3.65, *p* = .033; muscimol significantly lower than saline on Day 14 *t*(26) = −2.66, *p* = .013, but not Day 15 *t*(26) = −1.89, *p* = .071, or Day 16 *t*(26) = −1.35, *p* = .188). However, it is likely that this group difference in extinction reflects the pre-existing differences in responding at the end of stage 3. Overall, the pattern of data suggests that muscimol inactivation of OFC disrupts both learning and behavioural expression of simple Pavlovian cue-outcome associations.

#### OFC inactivation does not disrupt the motivation to consume food reward

The significant suppression of responding to all cues following OFC inactivation observed in stage 2 (Figure 2B and 3B) may have been a consequence of reduced motivation to consume the food reward. This explanation is unlikely given the absence of uneaten rewards following sessions in stage 2, however a more direct test of this explanation was necessary to rule out the possibility that the rewards were not eaten towards the end of the session when muscimol was no longer effective. Therefore, a consumption test was conducted within the test chambers with all animals being tested 10 mins after an infusion to ensure that the muscimol was maximally effective. Prior to the consumption test one muscimol and two saline group rats lost their cannula patency and were excluded from testing (saline n = 11, muscimol n = 14). All animals consumed all pellets by the end of the 30-minute session on both days, regardless of infusion group. Similarly, there was no evidence that muscimol infusion deferentially affected the vigour or frequency of magazine approach for freely available reward (Supplemental Figure S3; no significant effect or interactions with Group, all *F* < 0.94, *p* > .343).

### Experiment 2: OFC inactivation does not disrupt Pavlovian extinction learning by impairing the acquisition of conditioned inhibition

#### Histology

Cannulae placements are depicted in Figure 1C (representative image shown in Supplemental Figure S1B). All cannulae tips were located within LO or DLO. Final group numbers were saline (n = 12) and muscimol (n = 12).

#### Baseline responding

Rates of baseline responding did not significantly differ between groups during any of the testing phases and justified the analysis of CS-PreCS difference scores as measures of discriminative responding to the cues in consequent analyses. Briefly, one-way Group or Group x Day mixed ANOVAs were run separately for each stage of testing to assess the effects of Group. During stage 2 feature negative training (main effect of Group *F*(1, 22) = 2.55, *p* = .12; Group x Day interaction *F*(3, 66) = 1.72, *p* = .17), stage 3 testing (Group *F*(1, 22) = 3.32, *p* = .08), during stage 1 acquisition, summation and retardation tests all Group and Group x Day interactions failed to reach significance (all *F* < 1.72, *p* > .17).

#### Stage 1: Acquisition (days 1-9)

Acquisition of discriminative responding to cues A, B and C did not differ between (infusion) groups across stage 1 of acquisition (Supplemental Figure S4; significant main effect of Day *F*(8,176) = 26.07, *p* < .001, all remaining effects *F* < 0.83, *p* > .646; See Supplemental Figure S6 for responding to Cue Z).

#### Stage 2: OFC inactivation enhances within- but disrupts between session Pavlovian extinction (days 10-13)

Extinction of cue C following infusions in stage 2 allowed for a replication of the findings of (Panayi & Killcross, 2014) that OFC inactivation disrupts between- but not within-session extinction (Figure 4A). Extinction to cue A in compound with cue X was designed to test whether OFC inactivation impairs Pavlovian extinction by disrupting the formation of conditioned inhibition that may form during extinction (Delamater, 2004; Rescorla, 1969). Overall, in both groups extinction to AX- was significantly slower than to C- (significant Cue*Day interaction *F*(3,66) = 2.93, *p* = .040, but no interactions between Cue and Group, Day, or trial Block *F* < 2.30, *p* > .112; both cues showed significant linear decreases in responding over days, but this was faster for C-, *t*(22) = −5.76, *p* < .001, than AX-, *t*(22) = −2.79, *p* = .032). Reduced responding to the compound AX-compared to C- is consistent with external inhibition or generalisation decrement accounts of the novel presence of cue X suppressing responding.

**Figure 4.**
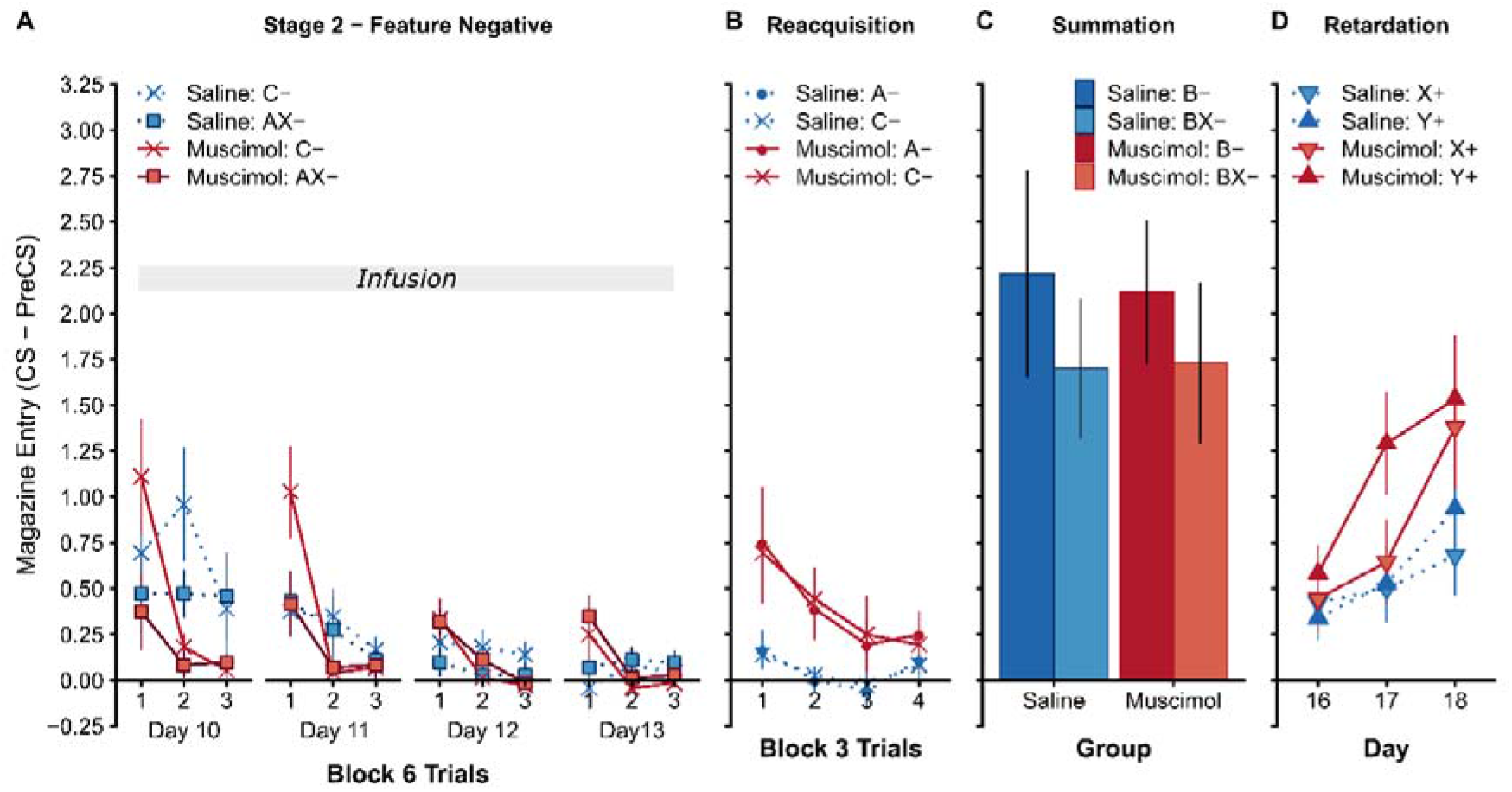
OFC inactivation disrupts between-session extinction and enhances within-session extinction and does not depend on the acquisition of conditioned inhibition. Experiment 2 **(A)** Extinction of AX- and C-during stage 2 of the conditioned inhibition procedure depicted in blocks of 6 trials within each session. Following saline infusions, responding to AX- and C-declined within- and between-sessions whereas muscimol infusions into LO impaired the retention of extinction between-session extinction. **(B)** Test of the responding to A- and C-drug-free depicted in blocks of 3 trials within the session. Responding in the muscimol group was significantly higher than the saline group but responding did not differ between cues. **(E)** A summation test revealed lower responding to BX-than B- in both the saline and muscimol groups. **(F)** A retardation test revealed similar rats of acquisition to X+ and control cue Y+ in muscimol and saline group. Rates of discriminative magazine responding presented as CS-PreCS difference scores in 10s. Error bars depict +SEM.

Extinction between-sessions was significantly impaired in the muscimol group (Group*Day interaction *F*(3,66) = 2.93, *p* = .040, main effect of Day *F*(3,66) = 17.65, *p* < .001). This effect was confirmed by a planned analysis looking only at the first block within each day (main effect of Group *F*(1,22) = 4.46, *p* = .046, and Day *F*(3,66) = 10.19, *p*< .001) as a measure of between-session extinction retention (i.e. a savings test; Panayi & Killcross, 2014). Muscimol inactivation significantly impaired but did not completely block between-session extinction.

In contrast to impaired between-session extinction, muscimol significantly enhanced within-session extinction compared to the saline group (Day*Block interaction *F*(6,132) = 2.52, *p* = .024, main effect of Block *F*(2,44) = 24.40, *p* < .001). Overall, muscimol responding was greater than Saline in Block 1 (*t*(22) = 2.11, *p* = .046), but significantly lower than saline in Block 2 (*t*(22) = −3.35, *p* = .003), and Block 3 (*t*(22) = −2.53, *p* = .019).

#### Stage 3 Extinction test (Day 14)

Drug free tests of A- and C-revealed that the muscimol group responded significantly higher than the saline group (Figure 4B; main effect of Group *F*(1,22) = 16.02, *p* = .001). Impaired retention of extinction to cue C in the muscimol group when tested drug free successfully replicates the findings of (Panayi & Killcross, 2014) showing that OFC inactivation disrupts extinction learning. Surprisingly, there was no evidence that compound extinction of cue A with cue X had “protected” cue A from extinction relative to cue C, in fact the mean responding to both cues were identical in both groups (main effect of Cue *F*(1,22) = 0.00, *p>* .999, no interactions with Group, Day, and trial Block *F* < 0.13, *p* > .944).

#### Stage 4 Summation test (Day 15)

Responding to compound BX- was lower than to cue B-alone in both groups at test (Figure 4C; significant main effect of Cue *F*(1,22) = 4.67, *p* = .042, but no effect of Group *F*(1,22) = 0.00, *p* = .956, or Group*Cue interaction *F*(1,22) = 0.10, *p* = .752). Therefore, the summation test provided evidence of conditioned inhibition to cue X in both groups.

#### Stage 5 Retardation test (Days 16-18)

Responding during the retardation test suggested that the rate of acquisition to cue Y was greater than cue X in the muscimol group but not the saline group (Figure 4D). However, this observation was not fully supported statistically (only a significant main effect of Day*F*(2,44) = 10.87, *p* < .001, all remaining effects *F* < 2.68, *p* > .116). Taken together, the failure to pass both the summation and retardation test indicates that there is insufficient evidence of conditioned inhibition to cue X (consistent with reports that this effect is not strong, Rescorla, 1979). This suggests that conditioned inhibition does not significantly contribute to the extinction procedure employed here and previously (Panayi & Killcross, 2014).

#### OFC inactivation does not impair spontaneous locomotor activity

One possible account of the enhanced within-session extinction observed in the muscimol group in Stage 2 is that muscimol infusions generally suppressed locomotor activity and exploration. A novel context locomotor assay was performed in these animals, again following infusion of either saline or muscimol which revealed no difference in the total levels or time course of exploratory activity (Supplemental Figure S5; significant main effect of Block *F*(11,242) = 61.88, *p* < .001, but no effect of Group *F*(1,22) = 0.00, *p* = .955, or Group*Block interaction *F*(11,242) = 0.54, *p* = .873) or orienting behaviour between groups (significant main effect of Block *F*(11,242) = 50.97, *p* < .001, but no effect of Group *F*(1,22) = 0.05, *p* = .821, or Group*Block interaction *F*(11,242) = 0.78, *p* = .664). This rules out non-specific behavioural accounts of the effects of intra-OFC muscimol inactivation.

## Discussion

Our results reveal a selective role for the OFC in inhibitory behavioural control, which partially supports the traditional hypothesis of OFC function as a source of inhibitory control over well-established behavioural responses. However, despite the abolition of selective inhibitory behavioural control following OFC inactivation, OFC function was not necessary for the underlying learning of an inhibitory association, as assessed by summation and retardation tests. This suggests that the learning and subsequent expression of conditioned inhibition are neurally dissociable. However, this finding is not consistent with latent state models of OFC function which predict intact behavioural control and learning when changes in reinforcement are cued by external explicit cues (i.e. cue X). This dissociation between learning and behaviour following lateral OFC inactivation in rodents is broadly consistent with recent findings in non-human primates (Murray, Moylan, Saleem, Basile, & Turchi, 2015) and in humans (Bechara, Damasio, Damasio, & Anderson, 1994; Fellows, 2011), which suggest that updating expected outcome values and translating this knowledge into behaviour can be dissociated within OFC subregions.

The fundamental impairment in behavioural control following OFC inactivation in the present studies cannot simply be attributed to failed inhibitory control as there were multiple instances of enhanced behavioural inhibition following OFC inactivation. Firstly, OFC inactivation in experiment 1 disrupted the behavioural discrimination by suppressing responding to both the rewarded cue (A+) and the non-rewarded compound (AX-). Secondly, this impairment in increasing responding was also observed with the acquisition of responding to control cue Z+ in experiment 1. Finally, experiment 2 found evidence of enhanced behavioural inhibition within extinction sessions following OFC inactivation. Thus, an account of the OFC as the neural locus of learning inhibitory associations, or even general inhibitory behavioural control, does not adequately describe the bidirectional disruption of behavioural control observed in the present studies.

This conclusion is consistent with population and single-unit neuronal activity recordings in the rodent OFC. For example, in a stop-signal task that requires the use of cues to guide correct behaviour (Bryden & Roesch, 2015), OFC activity was sensitive to the direction of responding and this activity was enhanced when the observed behaviour required suppression of an alternative response. This suggests that the OFC is involved in inhibitory behavioural tasks because it plays a role in guiding and boosting behavioural control of correct/chosen responses rather than the inhibition of incorrect responses. Indeed, a number of electrophysiological recording and lesion studies in rodents (Lucantonio et al., 2015; Morrison, Saez, Lau, & Salzman, 2011; Roesch, Calu, Esber, & Schoenbaum, 2010; van Wingerden et al., 2010) and primates (Chau et al., 2015; Murray et al., 2015; West, DesJardin, Gale, & Malkova, 2011) suggest that OFC activity tracks the expected value of cues used to guide behaviour. Therefore, situations in which disruption of OFC function impairs behaviour are likely to indicate deficits in selecting or increasing optimal behaviour based on their current motivational value within the array of possible behaviours. This account would explain deficits in inhibitory control following OFC damage as deficits in resolving response competition, and would also account for reports that the OFC is only necessary for modifying established behaviours rather than establishing control of de novo behaviours (Murray et al., 2007).

### Outcome expectancy guiding behaviour

A number of findings in experiment 1 suggest that OFC inactivation disrupted behavioural control rather than affecting learning *per se*. OFC inactivation disrupted behavioural control during the A+/AX-discrimination, but did not disrupt learning about the inhibitory properties of cue X. Similarly, OFC inactivation suppressed acquisition to control cue Z. While this effect persisted when OFC was subsequently functional, it also affected control cue B which was not presented during OFC inactivation. These findings are consistent with the representation of the value of expected outcomes in the OFC. Specifically, in Pavlovian cue-outcome conditioning the OFC is necessary for using the current motivational value of predicted outcome to guide conditioned behaviour (Gallagher et al., 1999; Pickens, Saddoris, Gallagher, & Holland, 2005; Pickens et al., 2003). In the absence of this information, behavioural control is likely to be guided by direct stimulus-response associations that may form during learning (Delamater, 2007; Hall, 2002; Holland & Straub, 1979; Killcross & Blundell, 2002). Thus, in simple cue-outcome learning to the control cue Z, the suppression in response acquisition following OFC inactivation may represent the loss of information about the current value of the reward that normally boosts responding, and a reliance on a slower stimulus-response memory system. Similarly, during the A+/AX-feature negative discrimination the loss of information about the current value of the reward would not be available to boost responding to A+. However, an inhibitory association between X and responding (or an excitatory connection between X and a specific representation of no-outcome (Delamater, 2004; Konorski, 1967) could still form with only a stimulus-response system. Therefore, while we provide direct evidence that learning of conditioned inhibition is not disrupted by OFC inactivation, further evidence is required to test whether behaviour, but not learning, is disrupted during simple cue-outcome learning.

One alternative account of OFC function that is consistent with the impaired acquisition to cue Z in experiment 1 is an impairment in correct credit assignment (Walton et al., 2011). While the reinforcement contingency of cue Z remained stable when muscimol infusions began, a non-reinforced cue was also introduced at this stage (AX-). Therefore, it is possible that OFC inactivation led to incorrect assignment of non-reinforcement to cue Z. However, it is not clear why this failure of credit assignment would not also disrupt the learning of conditioned inhibition to cue X.

### Outcome expectancy for learning

We have previously shown that the OFC is necessary for learning about cues that no longer signal reward in extinction procedures (Panayi & Killcross, 2014). The present findings (experiment 2) extend this to show that the specific extinction procedure parameters employed do not generate significant conditioned inhibition to other cues, and that OFC is not necessary for inhibitory cue-outcome learning in general. These findings are problematic for a strict view in which the OFC is the neural locus of outcome expectancy information that is involved in computing prediction errors necessary for learning. From a theoretical perspective, outcome expectancy information is a necessary component for prediction error signals which are critical to learning in associative learning (Mackintosh, 1975; Pearce & Hall, 1980; Rescorla & Wagner, 1972) and reinforcement learning theories (Sutton & Barto, 1998). Therefore, a straightforward prediction of the view that the OFC represents outcome expectancy signals that inform prediction errors is that disrupting OFC function should disrupt mid-brain prediction error signalling and fundamentally disrupt learning. However, this is refuted by a number of reports that OFC lesions do not disrupt initial task learning (Boulougouris, Dalley, & Robbins, 2007; Chang, 2014; Gallagher et al., 1999; Ostlund & Balleine, 2007; Scarlet, Delamater, Campese, Fein, & Wheeler, 2012), and do not disrupt putative reward prediction errors signals in the VTA in a manner consistent with the loss of outcome expectancy information (Takahashi et al., 2011).

To account for intact initial learning of tasks following OFC damage, a number of theories of OFC function (Delamater, 2007; Rudebeck & Murray, 2011; Schoenbaum et al., 2009) appeal to the distinction between learning about different aspects of rewards such their sensory specific properties (e.g. taste, shape, colour, location etc.) and their general motivational and rewarding properties (Dickinson & Dearing, 1979; Wagner & Brandon, 1989). The OFC is argued to represent outcome expectancy information about the sensory-specific properties of outcomes, leaving learning about the general properties of rewards intact and capable of supporting acquisition in the absence of OFC function. However, when correct task performance depends on representing the specific properties of the outcome, such as when a specific outcome changes value, then OFC function is necessary for correct performance (Gallagher et al., 1999; Murray et al., 2015; West et al., 2011). However, this explanation does not account for deficits in extinction learning in which the specific properties of the outcome are not relevant to task performance (Panayi & Killcross, 2014). Instead, Panayi and Killcross (2014) hypothesised that the disruption of expected outcome value information reduced the motivational significance of the extinction session in which no rewards were delivered. Consequently, responding at the start of the session may be driven by stimulus-response associations, but the lack of motivationally significant events in the chamber may facilitate rapid habituation to the cue and the context, protecting the responding from substantial extinction learning. This is supported by evidence of rapid within-session extinction found by Panayi & Killcross (2014) which was replicated in experiment 2, and has been observed in earlier reports (see figure 3E in Keiflin, Reese, Woods, & Janak, 2013). Indeed, the increased number of non-rewarded trial types in experiment 2 may account for the clearer evidence of rapid within-session extinction compared to that originally reported by (Panayi & Killcross, 2014).

### Creation and representation of latent states

A more recent theoretical approach has been to link OFC function with model-based reinforcement learning (Keiflin et al., 2013; McDannald et al., 2011, 2012; Takahashi et al., 2011; Wilson et al., 2014). Similar to the sensory-specific/general-motivational distinction, reinforcement learning distinguishes between model-based, sensory rich learning about task structure and specific reward properties, and model-free, general learning about non-specific average reward rates associated with cues or actions (Daw, Niv, & Dayan, 2005). The possibility that the OFC is involved in model-based learning extends the scope of OFC function to include representing task structure (Behrens et al., 2018; Niv, 2019) as well as specific properties of outcomes.

Within this framework, Wilson et al. (2014) modelled deficits in extinction following OFC damage as a deficit in representing a change in the task based on latent information such as associative history rather than explicit environmental cues. With an intact OFC, subjects can detect that the rate of reinforcement has changed during an extinction session and create a new task state representation in which new context-specific inhibitory learning can be acquired. In the absence of a functioning OFC, the subject cannot use internal information about the associative history of the cue to generate this new latent task state, and instead must directly update the original acquisition memory. Therefore, extinction learning in the absence of OFC function is modelled as erasure of the original learning rather than new context specific inhibitory learning. One prediction of this latent state account is that OFC damage should spare performance in tasks where changes in task structure are explicitly signalled with physical cues. Therefore, conditioned inhibition should not be disrupted by OFC inactivation since changes in reinforcement in the task are cued by the physical presence of cue X.

As predicted, OFC inactivation did not prevent learning of conditioned inhibition to cue X in Experiment 1. However, OFC inactivation significantly disrupted discriminative behavioural control during the learning stage (stage 2) which is not consistent with the model’s prediction. One possible reason for the discrepancy between the present findings and the model proposed by Wilson et al (2014) is the functional heterogeneity amongst the subregions of both the rodent and primate OFC (Chau et al., 2015; Izquierdo, 2017; Panayi & Killcross, 2018; Parkes et al., 2018; Rudebeck & Murray, 2011). In particular, the model based simulations of deficits during extinction are modelled on lever responding primate data from (Butter, Mishkin, & Rosvold, 1963) in which ablations of the entire orbital surface were conducted. Therefore it is likely that the model accounts for a range of OFC functions across multiple orbital regions (Bradfield, Dezfouli, van Holstein, Chieng, & Balleine, 2015; Sharpe, Wikenheiser, Niv, & Schoenbaum, 2015).

There are a number of diverse regions that have uniformly been considered as OFC regions (Price, 2006a; Roesch & Schoenbaum, 2006), however there is mounting evidence that these subregions are functionally heterogeneous in rodents and primates (Balleine et al., 2011; Mar, Walker, Theobald, Eagle, & Robbins, 2011; Rudebeck & Murray, 2011; Walton et al., 2015). In the present experiments cannula tips were restricted to the anterior portion of the lateral OFC. This is in contrast to the majority of rodent OFC studies that target the posterior portion of the lateral OFC with cannulae and neural recording probes, or excitotoxic lesion studies that can encompass lateral OFC, ventral OFC and agranular insular cortex (Chudasama & Robbins, 2003; Gallagher et al., 1999; Pickens et al., 2005; Scarlet et al., 2012). Rodent ventral and lateral OFC are functionally dissociable from the medial OFC (Mar et al., 2011), and ventral OFC appears dissociable to lateral OFC (Balleine et al., 2011; Corwin, Fussinger, Meyer, King, & Reep, 1994). There is even evidence of a dissociation of function across the anterior-posterior axis of the lateral OFC (Panayi & Killcross, 2018).Furthermore, these orbital subregions have distinct patterns of connectivity within the medio-dorsal thalamus, amygdala, and striatum in both rodents and monkeys (Groenewegen & Uylings, 2000; Hoover & Vertes, 2011; McDonald, 1998; Price, 2006b; Schilman et al., 2008). Functional heterogeneity in primates has also been shown between Walker’s areas 11, 12 and 13 (Murray et al., 2015; Noonan et al., 2010; Rudebeck & Murray, 2011; Walton, Behrens, Buckley, Rudebeck, & Rushworth, 2010). It is therefore important to start discriminating between OFC subregions when characterising the function of the OFC and attempting to establish homologous regions between species and clarify inconsistent findings.

### Summary

These findings suggest a role for the rodent lateral OFC in the online behavioural control but not the underlying associative learning of conditioned inhibition. These findings are consistent with a role for the OFC in guiding current behaviour based on the expected value of predicted outcomes. Dysfunction of the OFC is consistently implicated in psychological disorders of behavioural control such as obsessive compulsive disorder (Burguiere, Monteiro, Feng, & Graybiel, 2013; Graybiel & Rauch, 2000). These are usually thought of as disorders of inappropriate response inhibition, however the present findings suggest that they may also be considered disorders of appropriate response selection.

## Acknowledgements

We gratefully acknowledge Fred Westbrook, Nathan Holmes, David Bannerman, Mark Walton, and Geoffrey Schoenbaum for their invaluable feedback. Research supported by grants awarded to Simon Killcross from the Australian Research Council (ARC Discovery Grant DP0989027 and DP120103564).

## Data availability

Data available online at: https://osf.io/jt958/?view_only=5effc55b5cb5408bbf305f8c29755ecb

