## Supplementary Figures for "Orbitofrontal Cortex is necessary for the behavioural expression, but not learning, of Pavlovian conditioned inhibition"

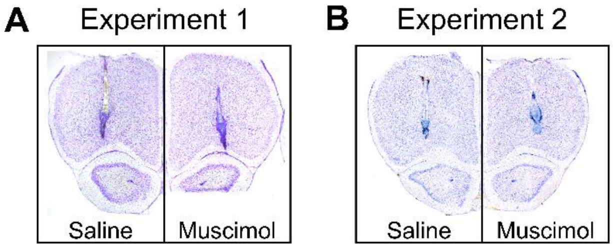


**Supplemental Figure S1.** Representative photomicrographs of Nissl stained coronal sections depicting cannulae placement in the lateral OFC in the saline (left) and muscimol (right) groups in **(A)** Experiment 1, and **(B)** Experiment 2. Placements estimated at 4.20 mm relative to bregma (Paxinos and Watson, 1997).

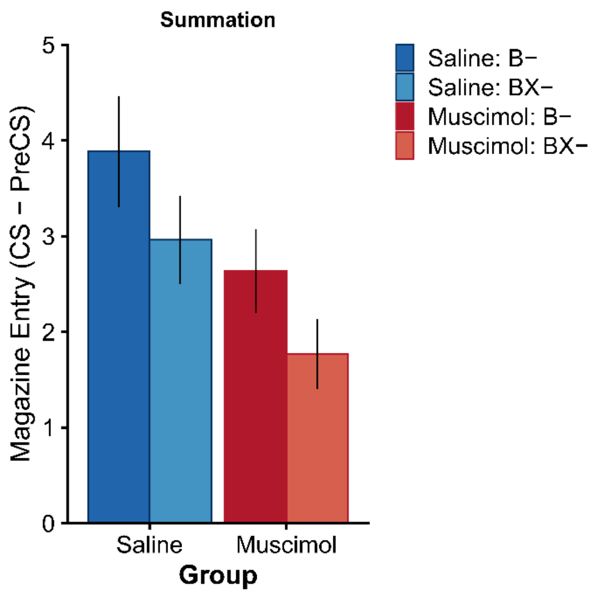


**Supplemental Figure S2.** The first block of two trials from the summation test in Experiment 1 (full test data shown in Figure 2D). Rates of discriminative magazine responding in presented as CS-PreCS difference scores in 10s. Error bars depict ±SEM.

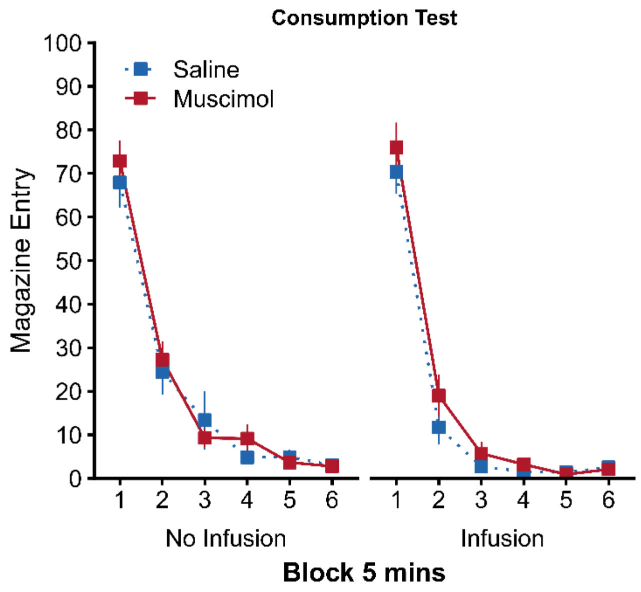


**Supplemental Figure S3.** OFC inactivation does not affect appetite or vigour of reward approach when rewards are freely available. Magazine frequency in 5-minute blocks following a dummy infusion (No Infusion; Left) and intra- OFC drug infusion (Infusion; Right). All animals ate the 40 reward pellets that were placed in the magazine and freely available from the start of the 30-minute test sessions.

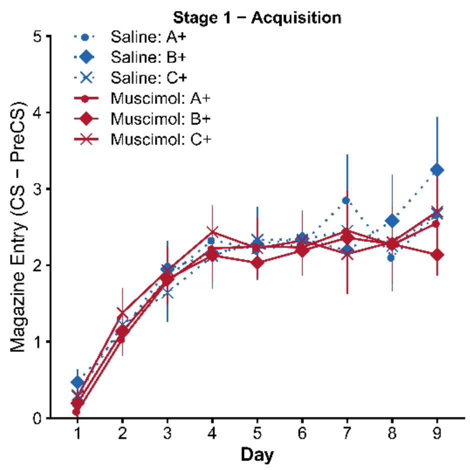


**Supplemental Figure S4.** Acquisition of conditioned responding to cues A+, B+, and C+ before (days 1-6) and after surgery and post-operative recovery (days 7-9). Rates of discriminative magazine responding presented as CS-PreCS difference scores in 10s. Error bars depict +SEM.

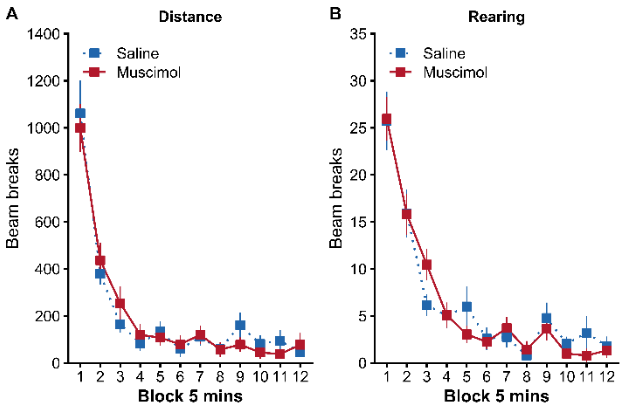


**Supplemental Figure S5.** OFC inactivation does not disrupt locomotor activity and novelty exploration in a locomotor assay. **(A)** Total distance travelled in blocks of 5 minutes as measured by the total number of infra-red beam breaks. **(B)** Frequency of rearing behaviour travelled in blocks of 5 minutes as measured by the total number of infra-red beam breaks located 14 cm above the floor. Error bars depict +SEM.

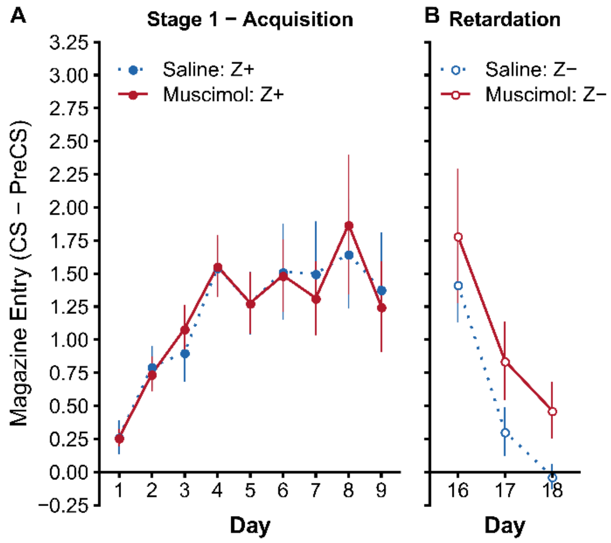


**Supplemental Figure S6.** Responding to control cue Z in Experiment 2 during **(A)** Stage 1 acquisition, and (B) non-reinforcement during the retardation test. Rats in both groups acquired responding to Z+ in Stage 1 at a similar rate (significant main effect of Day $F(8{,}176) = 8.80$, $p < .001$, but no effect of Group $F(1{,}22) = 0.00$, $p = .997$, or Group*Day interaction $F(8{,}176) = 0.18$, $p = .994$). During the retardation test, responding to non-reinforced cue Z- also decreased at a similar rate in both groups (significant main effect of Day $F(2{,}44) = 20.78$, $p < .001$, but no effect of Group $F(1{,}22) = 2.07$, $p = .164$, or Group*Day interaction $F(2{,}44) = 0.08$, $p = .925$). Rates of discriminative magazine responding presented as CS-PreCS difference scores in 10s. Error bars depict +SEM.
